# Removal of Hg and Cu from raw gold-mine tailings in microalgae- and bacteria-augmented microbial fuel cells: Effects of bio-augmentation, diurnal cycling and operational faults

**DOI:** 10.64898/2026.09.02.748966

**Authors:** Joana Iza, Jennifer Cuadrado, Lourdes García-Rodríguez, Celso Recalde, Petteri Nurmi, Agustin Zuniga

## Abstract

Artisanal gold-mining tailings carry dissolved mercury, copper and lead at concentrations requiring treatment before discharge. Bench-scale microbial fuel cells (MFCs) offer passive treatment of such streams, but exhibit operational variability that obscures whether removal is biologically driven. This study applies an integrated diagnostic framework to two bench-scale systems treating tailings from two artisanal gold-mining districts in Ecuador. System 1 evaluated *Micractinium inermum* bio-augmentation, while System 2 compared *Psychrobacter alimentarius*- and *Trichococcus patagoniensis*-dominated anodic consortia. The framework combines paired time-series statistical modeling, an adaptive percentile-floor change-point detector, equivalent-circuit modeling, and baseline-corrected XRD/FTIR spectroscopy. In System 1, treatment reduced mercury from 0.042 to 0.0027–0.0148 mg L*^−^*^1^ and copper from 2.79 to 0.130–1.837 mg L*^−^*^1^. Against the applicable discharge limits, copper complied in six of eight reactor–phase combinations, whereas mercury exceeded its limit throughout, identifying it as the limiting contaminant. Removal was governed by system-level physicochemical processes rather than algal-specific biocatalysis, with the non-algal control matching or exceeding the augmented reactors. Paired per-day testing showed that algal bio-augmentation conferred no voltage advantage under stable operation (+0.17%) but a +27.54% gain under diurnal perturbation, periodicity analysis attributed these shifts to the 24-h chamber photoperiod. An adaptive change-point detector distinguished recoverable excursions from terminal structural collapse in both systems, and equivalent-circuit fitting attributed System 2 consortium power difference to internal resistance and open-circuit voltage. The framework separates biological treatment effects from mechanical failure, supporting automated health monitoring in bio-electrochemical wastewater treatment.

## 1. Introduction

Artisanal and small-scale gold mining generates effluents laden with dissolved mercury, copper and lead derived from ore-processing tailings, constituting a persistent environmental and public-health liability at the operations where the contamination originates [1, 2, 3]. Conventional treatment of these streams is capital- and energy-intensive, which restricts deployment where it is most needed. Microbial fuel cells (MFCs) have been proposed as a dual-function technology for such streams, generating electrical output while immobilizing or transforming dissolved metals through anodic bio-sorption, bio-reduction and precipitation [4, 5, 6, 7, 8]. Reported metal-removal performance is dominated by engineered dual-chamber configurations operating on synthetic media or catholytes, which achieve recoveries exceeding 99% [9, 10, 11, 12]. Such architectures nonetheless carry high internal resistance, ion-exchange membrane fouling and capital costs that render them impractical for continuous treatment of raw, unbuffered tailings [13, 14]. Single-chamber and sediment-type designs reduce both cost and membrane-associated ohmic losses, but are more variable and subject to uncharacterized background attenuation [15, 16]. Few studies separate biological metal attenuation from abiotic sorption and mineral precipitation, leaving it unresolved whether removal is attributable to a specific biocatalyst or to the reactor environment as a whole [17, 18].

Photosynthetic microalgae such as *Micractinium inermum* have been co-cultivated in bio-electrochemical systems to provide in situ pH buffering and cathodic oxygenation without mechanical aeration or synthetic electron acceptors [19, 20, 21]. Microalgae-assisted and photosynthetic bio-voltaic cells operating on agricultural or domestic wastewater report power densities of 34-59 mW m*^−^*^2^ [22, 23, 24]. Two analytical constraints recur across this literature. Single-reactor-per-condition designs remain widespread [10, 11], leaving reported gains exposed to temporal pseudoreplication and reactor-to-reactor variance [25, 26], and algal–bacterial studies rarely isolate algae-specific enhancement from system-level sorption and bio mineralization [23].

Electrogenic output also depends on the composition and extracellular electron transfer efficiency of anodic biofilm. Comparisons between psychrotolerant taxa such as *Psychrobacter alimentarius* [27] and electrochemically active fermentative genera such as *Trichococcus patagoniensis* [28] show that community structure modulates charge-transfer kinetics, internal resistance (*R*_int_) and open-circuit voltage (*V*_oc_). Polarization sweeps and impedance spectroscopy are standard characterization tools, but single-point metrics such as maximum-power-point resistance are distorted by activation losses and concentration polarization near the sweep extremes [29, 30, 31, 32, 33]. Reconciling an ohmic-region equivalent-circuit fit against the single-point estimate offers a non-invasive indication of where those losses arise [34, 35].

Scale-up is further limited by operational instability, electrode bio-fouling, separator breaches and voltage reversal [36, 37, 38]. Performance is commonly assessed using time-averaged statistics, which attenuate high-frequency fluctuations and obscure early failure signals. Data-driven change-point and anomaly-detection methods are well established in industrial process control [39, 40], yet tuning-free operando tools adapted to bio-electrochemical time series remain scarce, and existing approaches do not distinguish sudden structural failure from progressive baseline drift [41, 42]. Without such tooling, stochastic hardware failures can be easily misattributed to biological treatment effects, potentially leading to incorrect conclusions about the underlying mechanisms governing electrochemical behavior.

This study addresses these gaps by applying an integrated diagnostic framework to two bench-scale MFC systems treating tailings from two artisanal gold-mining districts in Ecuador: Nambija (Zamora Chinchipe) for System 1 and Zaruma–Portovelo (El Oro) for System 2. The first comprises four single-chamber reactors, operated in two phases separated by physical reconstruction of all cells, comparing *M. inermum* against a non-algal control under continuous open-circuit monitoring. The second comprises four two-electrode cells comparing *P. alimentarius*- and *T. patagoniensis*-dominated anodic consortia. The framework couples paired per-day statistical testing, adaptive change-point detection, periodicity analysis, equivalent-circuit fitting, and baseline-corrected XRD and FTIR characterization.

The objectives of this study were to (i) quantify mercury and copper removal from raw gold-mine tailings and determine whether it is attributable to algal biocatalysis or to system-level processes, (ii) establish, using a paired per-day design, whether algal bio-augmentation confers a measurable electrical benefit and under which operating conditions, (iii) develop an adaptive change-point detector able to distinguish recoverable excursions from terminal structural failure in continuous voltage records, and evaluate it across two independent reactor configurations, (iv) resolve the power-density difference between anodic consortia into its internal-resistance and open-circuit-voltage components, and (v) characterize mineralogical and surface-chemical changes resulting from treatment relative to untreated tailings.

## 2. Materials and Methods

Two bench-scale MFC systems were analyzed using an analytical framework that bridges continuous process monitoring with physical characterization. Section 2.1 describes the MFC configurations that were analyzed and provides an overview of the monitored parameters. Section 2.2 presents the methodology used to analyze heavy metal remediation. Section 2.3 describes the computational diagnostic framework used to analyze and characterize faults according to their severity, while Sections 2.4 and 2.5 describe the electrochemical modeling and solid-phase spectroscopic methods used to diagnose the physical root causes of the computational triggers. Table 1 summarizes the experimental systems and analytical methods used on them.

**Table 1:** Summary of experimental setups and analytical methodologies.

| Parameter | System 1: Algal Bio-augmentation | System 2: Microbial Communities |
| --- | --- | --- |
| **Reactor Configuration** | 4 single-chamber MFCs (3 algal, 1 control) | 4 two-electrode MFCs (2 *P. alimentarius*, 2 *T. patagoniensis*) |
| **Duration** | 18.87 days (Phase 1: 9.68, Phase 2: 9.19) | 19–21 days |
| **Continuous Monitoring** | Cell voltage (5-second logging resolution) | Cell voltage (1-minute logging resolution) |
| **Target Application** | Heavy metal remediation (macroscopic scale) | Anodic biofilm characterization (microscopic scale) |
| **Analytical Methods** | Aqueous heavy metal analysis (Hg, Cu, Pb) | Polarization sweeps, XRD, FTIR spectroscopy |
| **Computational Analysis** | Change-point detection, hypothesis testing | Change-point detection |

### 2.1. Physical Reactor Configurations and Analytical Datasets

#### Algal Bio-augmentation Operational Setup (System 1)

The system comprised four single-chamber MFCs operated over two discrete phases: an initial operational period (Phase 1: 9.68 days) and a secondary period following cell reconstruction (Phase 2: 9.19 days), with a total operating time of 18.87 days, to evaluate algal bio-augmentation for gold-mine tailings remediation. The two phases were separated by a planned cell reconstruction interval for all SMFCs. To preserve the biological community across the transition, the anodic biofilm and algal inoculum were retained and transferred to the reconstructed cells. Three reactors were bio-augmented with the microalga *M. inermum* (SMFC1, SMFC2, and SMFC4), while one reactor operated as an unaugmented non-algal control (SMFC3). All reactors were housed in a single environmental chamber maintained at 23 *^∘^*C and illuminated with white LEDs on a 06:00–20:00 photoperiod at an incident intensity of 2300 lux, measured at the reactor surface (*≈*31 µmol photons/m^2^/s to 43 µmol photons/m^2^/s photosynthetically active radiation). The reactor vessels were transparent, so that chamber illumination reached the algal suspension directly, and were maintained in equivalent positions throughout both phases. Cell voltages were recorded continuously under open-circuit conditions (no external load applied) at 5-second temporal resolution.

#### Anodic Microbial Community Dynamics (System 2)

This system evaluated anodic microbial community performance across four two-electrode MFCs operated over 19–21 days with continuous 1-minute voltage logging and no acquisition interruption. Tailings for this system were collected at 10 cm depth from a gold-processing plant in the Zaruma–Portovelo mining district (El Oro Province, Ecuador, 1200 m a.s.l.) and transported to the laboratory under refrigeration, stored in the dark until use. Duplicate reactors were inoculated with either a *P. alimentarius*-dominated consortium (C1, C2) or a *T. patagoniensis*-dominated consortium (T1, T2). Quasi-steadystate electrochemical polarization sweeps were acquired across 21 discrete external load resistance steps ranging from 50 Ω to 2 MΩ, following standardized bio-electrochemical characterization protocols [31, 32].

### 2.2. Aqueous Heavy Metal Analysis

Tailings and process water for System 1 were collected from a gold-extraction plant in the Nambija Mining District, Zamora Chinchipe, Ecuador (4*^∘^*04*^′^*13*^′′^* S, 78*^∘^*45*^′^*04*^′′^* W, WGS 84), during the dry season. The influent tailings water contained 0.042 mg L*^−^*^1^ Hg, 2.79 mg L*^−^*^1^ Cu and 0.37 mg L*^−^*^1^ Pb, together with 3 mg L*^−^*^1^ nitrate and 55 mg L*^−^*^1^ sulfate, at pH 8.3. Aqueous remediation performance, comprising the depletion of heavy metal concentrations (Hg, Cu, and Pb), was monitored exclusively for the first test system across the two operational phases defined in Section 2.1 (Phase 1 and Phase 2). Metal removal kinetics were quantified by periodically extracting liquid aliquots (100 mL) from the reactors. Prior to analytical quantification, all samples were filtered through 0.45 µm membrane filters to remove suspended biomass and solid particulates, followed by acidification with trace-metal grade nitric acid (HNO_3_) to a pH < 2. This preservation step was necessary to ensure metal stabilization and prevent localized adsorption to the sample vial walls. Acidified aliquots were refrigerated, retained for analysis, and were not returned to the reactors. All four reactors were retained in the removal analysis for both phases. The Phase 2 separator failure in SMFC4 affected the electrical path but not the aqueous sampling, which was performed independently of cell voltage.

Dissolved metal concentrations were quantified using an atomic absorption spectrophotometer (AAS–Thermo Scientific^TM^ iCE^TM^ 3500), following APHA Standard Methods for water-quality analysis. Calibration was performed using certified single-element standard solutions (Inorganic Ventures) at concentrations of 5, 10, 20, 30, and 40 µg L*^−^*^1^ for Hg; 0.1, 0.2, 0.4, 1.0, and 2.0 mg L*^−^*^1^ for Cu; and 0.3, 0.6, 1.2, 1.8, and 3.0 mg L*^−^*^1^ for Pb, and analytical blanks (ultrapure water, NOVA Laboratory) were run systematically to ensure analytical accuracy. Mercury was determined by cold-vapour generation, while copper and lead were determined by flame atomisation on the same instrument. Influent concentrations were measured by an external laboratory and post-treatment residuals in-house. The validated lead detection limit was 0.3 mg L*^−^*^1^. All post-treatment lead concentrations were below this limit and are reported as <0.3 mg L*^−^*^1^, precluding calculation of lead removal efficiency. Residual copper concentrations (0.130–1.837 mg L*^−^*^1^) were within the calibration range. Four of eight residual mercury concentrations were below the lowest calibration standard (5 µg L*^−^*^1^) but produced absorbances well above the blank.

The overall heavy metal removal efficiency (*R_E_*) was evaluated for each phase using the following relationship:

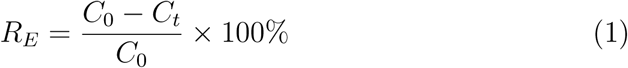

where *C*_0_ is the initial bulk metal concentration (mg L*^−^*^1^) at the start of the operational phase and *C_t_* is the residual metal concentration (mg L*^−^*^1^) measured at time *t*. Because the cells were reconstructed between phases (Section 2.1), *C*_0_ was re-established at the beginning of Phase 2. Residual concentrations were additionally compared against the discharge limits for freshwater bodies specified in Ecuadorian environmental legislation [43].

### 2.3. Computational and Analytical Diagnostic Framework

#### Voltage Time-Series Preprocessing and Statistical Design

Data quality control protocols were applied to continuous voltage streams to ensure physical fidelity prior to statistical evaluation. For System 1, raw voltage readings falling outside realistic bio-electrochemical boundaries (*V*_cell_ < 0.15 V or *V*_cell_ > 0.90 V), indicative of open-circuit disconnections or data acquisition saturation, were removed as sensor artifacts. The 0.15 V boundary lies within voltage range reached near reactor failure. The threshold excluded extreme low-voltage Phase 2 measurements from SMFC4 (5.4%) and SMFC3 (0.6%). No records measured *≤* 0 V. The reported minimum voltages are upper bounds, while reported collapse magnitudes are lower bounds.

Time was referenced to the start of each phase and analyses were restricted to individual phases (Section 2.1). Voltages were averaged into 30-minute intervals. Reactor-days were retained only if they contained *≥* 24 intervals and *≥* 80% valid data. Phase 1 comprised ten reactor-days for all four reactors, and Phase 2 nine reactor-days (eight for SMFC4). To address temporal autocorrelation and prevent pseudoreplication associated with high-frequency logging in single-reactor configurations, high-density voltage data were aggregated into daily mean cell voltages per reactor-day, with the daily standard deviation retained as a measure of operational stability. Paired *t*-tests, comparing the pooled algal treatment group against the non-algal control, were applied independently within each phase (Phase 1: *n* = 10, Phase 2: *n* = 9 paired reactor-days). Membrane failure excluded SMFC4 from the Phase 2 algal treatment pool. Moving-block bootstrap resampling provided 95% confidence intervals, accounting for day-to-day dependence without distributional assumptions. Phase contrasts were estimated using difference-in-differences, with confidence intervals obtained by block bootstrap.

#### Continuous Change-Point Detection and Degradation Ranking

To enable automated fault detection and distinguish physical operational degradation from biological fluctuations, an adaptive percentile-floor change-point algorithm was developed and applied independently within each operational phase. First, the continuous raw voltage time series *v*(*t*) is resampled into 30-minute discrete interval means and standard deviations (SDs). A 1-day rolling window (*W* = 48 intervals) is then applied to compute the moving temporal standard deviation *σ*(*t*) = std*{v*(*τ*): *τ ∈* [*t − W*, *t*]*}*. Windows containing fewer than 46 of their 48 constituent intervals were discarded. Detected onsets were insensitive to this requirement across the range 44–47 intervals.

To account for inherent background noise and avoid manual threshold tuning, a baseline noise floor *F* is established individually for each reactor. This floor is set as the 25th percentile of its rolling SD across the phase:

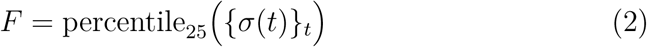

The algorithm then flags a degradation onset event (*t*_0_) the moment the rolling variance exceeds this noise floor and remains elevated. Specifically, the algorithm looks for periods where the variance is at least three times the noise floor and remains at that level for at least 70% of the subsequent six intervals (3-hour window):

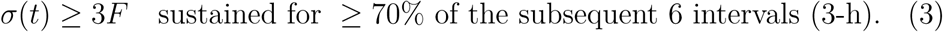

Once an event is flagged, the algorithm ranks fault severity by three complementary metrics applied in order of precedence: the onset day of the fault *t*_0_, the highest variance observed *σ*_max_ = max*_t_ σ*(*t*), and the fault severity ratio *S*:

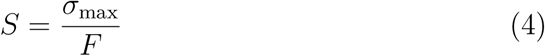

Finally, we calculate the voltage drop *D* to distinguish fatal (complete collapse) and transient (temporary instability) events:

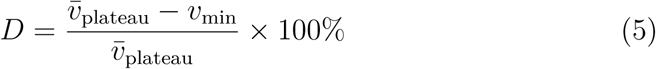

where *v̄*_plateau_ is the average voltage over the first three days of the phase, and *v*_min_ is the lowest voltage observed following the fault onset. These metrics provide insights into the nature of the degradation event rather than just providing a binary healthy/failed classification. The multi-metric formulation ranks reactors along a continuum of physical degradation modes, ranging from localized membrane breaches to progressive electrode fouling [41, 42]. The complete procedure is given as pseudocode in Supplementary Information (Algorithm S1).

To demonstrate applicability across distinct reactor configurations and operating conditions, the detector was applied to both test systems. To prevent the initial biological inoculation and acclimation phases from being misidentified as operational failures, a startup exclusion time (*t*_exclude_) was incorporated into the search window. This parameter was set to 1 day for each phase of the first system (the minimum required for the 1-day rolling window) and 6 days for the second system, corresponding to the empirical time required for anodic biofilm acclimation and stabilization. All subsequent algorithm operations, including noise floor computation, onset criteria, and severity metrics, remained identical across both test systems.

#### Diurnal Periodicity

To resolve sub-daily dynamics, the 30-minute data were binned into hourly deviation profiles, with uncertainty expressed as the cross-day standard error relative to amplitude. Periodicity was then quantified using Lomb–Scargle periodograms (4–72 h) on the detrended series to isolate the dominant period and normalized power per reactor.

### 2.4. Electrochemical Modeling and Internal Resistance Diagnostics

To analyze the operational limits and internal energy losses of the microbial communities, quasi-steady-state electrochemical polarization curves were acquired for the second test system across 21 discrete external load resistance steps (50 Ω to 2 MΩ). Current density (*J*, mA m*^−^*^2^) and power density (*P*, mW m*^−^*^2^) were normalized to the total electrode area (231.6 cm^2^: anode 168 cm^2^, cathode 63.6 cm^2^). Due to mass-transport limitations at near-short-circuit loads and activation losses near open circuit, polarization curves routinely exhibit non-linear features at sweep extremes [32, 29]. To isolate linear ohmic polarization behavior, each curve was segmented into three operational conditions: (i) a mass-transport-dominated region at low load resistance, (ii) an activation-dominated region near open circuit (500 kΩ–2 MΩ), and (iii) a linear ohmic-dominated mid-range (300 Ω to 100 kΩ, 13 sweep points).

Within the linear ohmic region, polarization behavior was modeled using standard equivalent-circuit formulation:

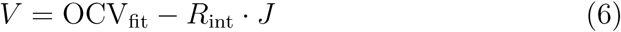

where *V* is cell voltage (V), OCV_fit_ is the extrapolated open-circuit voltage (V), *J* is current density (mA m*^−^*^2^), and *R*_int_ is the area-specific internal resistance (kΩ *·* m^2^). Parameters OCV_fit_ and *R*_int_ were determined via ordinary least-squares (OLS) regression across the 13 mid-range sweep points (Section 3.4).

To evaluate local non-ohmic losses at peak power production, a maximum-power-point (MPP) internal resistance (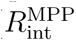) was independently computed:

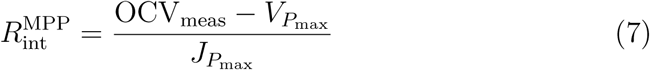

where OCV_meas_ is the measured open-circuit voltage at maximum load (*R*_ext_ = 2 MΩ), and *V_P_*_max_ and *J_P_*_max_ correspond to cell voltage and current density at maximum power output. Departure from ideal Thévenin impedance matching (*V_P_*_max_ /OCV = 0.5) [33, 34] was evaluated as a diagnostic index for operating point-specific overpotentials. The resulting area-specific internal resistances (0.0325 kΩ*·* m^2^ to 0.0601 kΩ*·* m^2^, equivalently 32.5 Ω*·* m^2^ to 60.1 Ω*·* m^2^) exceed values reported for mixed-culture bench-scale MFCs (0.009 Ω*·* m^2^ to 0.15 Ω*·* m^2^) [30] by two to three orders of magnitude.

### 2.5. Baseline-Corrected Spectroscopic Processing (XRD and FTIR)

Diffraction patterns and vibrational spectra were acquired to identify mineralogical transformations and biofilm-associated functional group shifts, and subsequently processed using baseline-corrected analytical protocols. For X-ray diffraction (XRD) data, patterns were acquired for untreated mine tailings and post-treatment anode samples from the *T. patagoniensis* (SMFC-T) and *P. alimentarius* (SMFC-C) reactors using Cu-K*α* radiation (2*θ* = 20*^∘^*–80*^∘^*, step size 0.01*^∘^*) [44]. Peak detection was then conducted following local baseline subtraction using a 200-point rolling-minimum filter, with candidate diffraction peaks flagged at a minimum relative prominence threshold of 2% of peak intensity. Primary mineral phases were matched against quartz reference diffraction profiles (SiO_2_) [45]. Because quartz is chemically and biologically inert under typical bio-electrochemical conditions, diffraction patterns were normalized relative to their respective quartz (101) reflection peak (2*θ ≈* 26.6*^∘^*), establishing an internal intensity standard for relative phase quantification. Non-quartz reflections were evaluated as candidate secondary mineral phases relative to untreated mine tailings baseline.

For Fourier-transform infrared (FTIR) spectroscopy, spectra were recorded across five anode and reference substrate samples using a JASCO FTIR spectrometer (wavenumber range 528–4003 cm*^−^*^1^, 902 channels) [44]. Transmittance spectra (*T*) were converted to integrated pseudo-absorbance (*A*) over the spectral range:

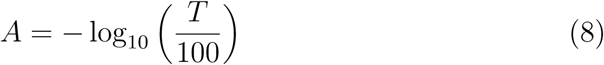

Biofilm-mineral surface interactions were evaluated by quantifying diagnostic absorption band in C–O/Si–O stretching region (1000–1010 cm*^−^*^1^) via local baseline depth and integrated band area across a fixed spectral window (950– 1150 cm*^−^*^1^) across all anode and reference samples, based on established vibrational band assignments for bio-electrochemical surface complexes [45].

#### Scanning Electron Microscopy

Surface morphology of the anode material was examined by scanning electron microscopy (JEOL JSM-IT100LA, 20.0 kV). Samples were air-dried and sputter-coated with gold prior to imaging. Micrographs were acquired for the anode material of all four System 2 reactors at magnifications ranging from 160*×* to 2200*×*. Images are reported qualitatively, to corroborate the FTIR band assignments; no quantitative image analysis was performed.

## 3. Results

### 3.1. Algal bio-augmentation confers a conditional voltage advantage

Unbuffered mine tailings introduce a high degree of operational variability and noise that can obscure biocatalytic performance, which the proposed statistical framework was developed to filter out. When applied to the algal bio-augmentation setup (System 1), continuous voltage monitoring demonstrates that the *M. inermum* effect varies with operating conditions (Figure 1 and Table 2). During Phase 1 (*n* = 10 paired reactor-days), the pooled algal bio-augmented group averaged a daily mean cell voltage of 0.5550(127) V, compared to 0.5541(100) V for the non-algal control. A paired *t*-test confirmed the +0.17% relative difference was not statistically significant (*p* = 0.866, 95% CI [*−*0.0120, +0.0179] V). During Phase 2 (*n* = 9 paired reactor-days), the algal group averaged 0.5778(197) V against 0.4530(687) V for the control. A paired *t*-test confirmed a +27.54% relative gain (124.8 mV) that was statistically significant (*p* = 0.0002, 95% CI [+0.0814, +0.1706] V). This phase dependence is consistent with cathodic oxygenation by the photosynthetic *M. inermum*. In single-chamber configurations, in situ oxygen evolution lowers cathodic overpotentials without mechanical aeration or synthetic electron acceptors, a benefit that remains latent while the system operates stably and becomes measurable only once it is perturbed.

**Figure 1:**
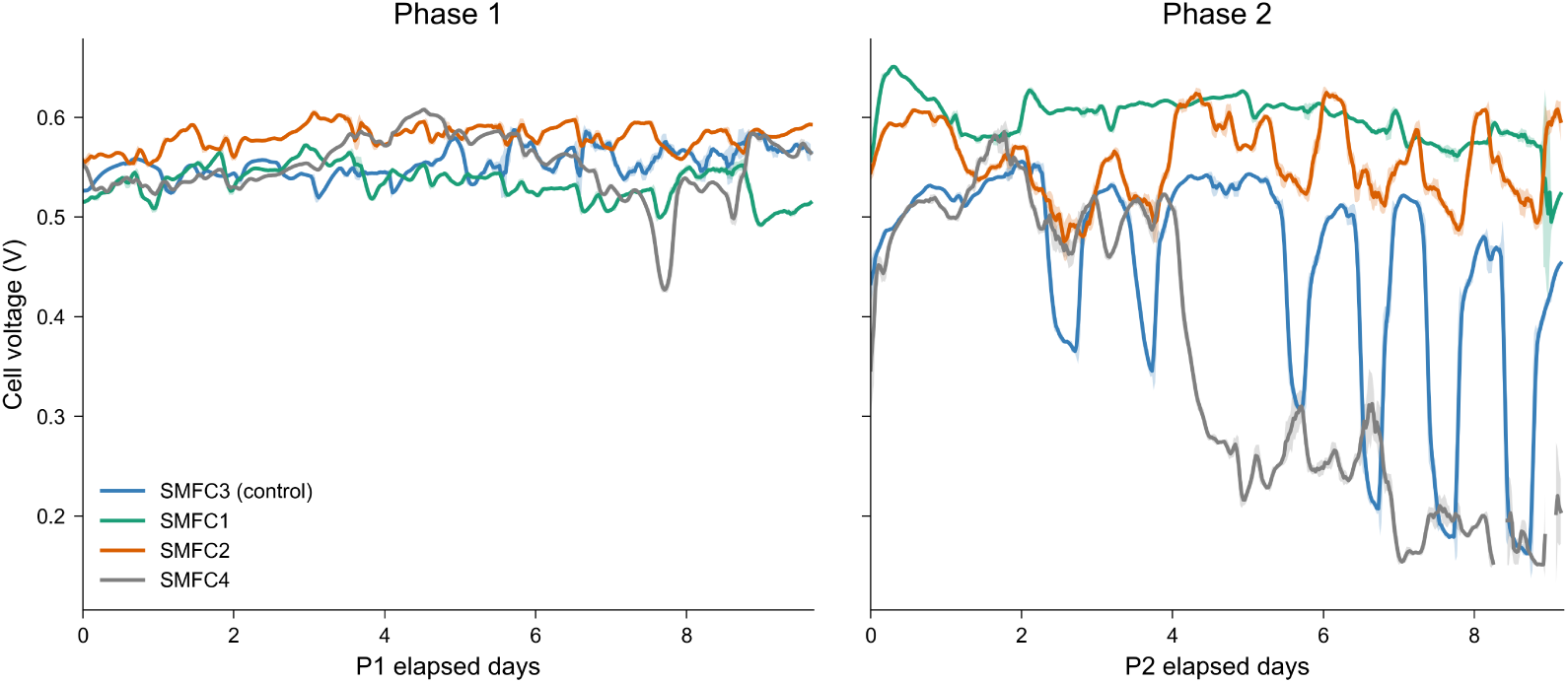
Continuous cell voltage profiles (2-h rolling mean *±* 1 SD) for algal (SMFC1, SMFC2, SMFC4) and control (SMFC3) reactors. Phase 1 (left) and Phase 2 (right). The control shows repeated Phase 2 voltage collapses, a pattern attenuated in algal reactors.

**Table 2:** Daily mean cell voltages by operational phase. Values across n-reactor-days (Phase 1: *n* = 10 for all SMFCs, Phase 2: *n* = 9 for three SMFCs and *n* = 8 for SMFC4).

| Reactor | Group | Phase 1 | Phase 2 |
| --- | --- | --- | --- |
| | | Mean ± SD, (V) | Mean ± SD, (V) |
| SMFC1 | Algae (*M. inermum*) | $0.5334 \pm 0.0146$ | $0.6002 \pm 0.0183$ |
| SMFC2 | Algae (*M. inermum*) | $0.5804 \pm 0.0086$ | $0.5554 \pm 0.0281$ |
| SMFC4 | Algae (*M. inermum*) | $0.5513 \pm 0.0273$ | $0.3804 \pm 0.1403$ |
| SMFC3 | Control (no biocatalyst) | $0.5541 \pm 0.0100$ | $0.4530 \pm 0.0687$ |
| **Pooled algae** |  | **$0.5550 \pm 0.0127$** | **$0.5778 \pm 0.0197$** |

Difference-in-differences quantified the between-phase shift at 0.1238 V (95% CI [+0.0731, +0.1673] V). Between phases the control mean fell from 0.5541 V to 0.4530 V and its mean within-day standard deviation increased 6.7-fold (0.0111 V to 0.0741 V), whereas the algal group rose from 0.5550 V to 0.5778 V with only a 1.7-fold change (0.0108 V to 0.0185 V). Across Phase 2 reactor-days the algal pool was less variable than the control by 0.0556 V (95% CI [*−*0.0891, *−*0.0273]). In Phase 1, reactor SMFC4 operated at 0.5513(273) V, consistent with the other bio-augmented reactors; retaining SMFC4 in the Phase 2 pool reduces the gain to +9.82% (*p* = 0.0062). Phase 1 also establishes an empirical baseline for reactor-to-reactor variability, with SMFC1 and SMFC2 deviating from the control by *−*3.72% (*p* = 0.019) and +4.75% (*p* = 0.0001). The Phase 2 differential exceeds this largest single-reactor deviation 5.8-fold, indicating that it cannot be attributed to inter-reactor construction variance alone. Taken together, algal bio-augmentation conferred no measurable advantage during stable operation but preserved cell potential under operational perturbation [46], indicating that its benefit in complex tailings matrices is conditional on the operating regime rather than intrinsic. The paired design with block-bootstrap intervals isolates this effect from day-to-day temporal autocorrelation, which a conventional summary-statistic comparison would absorb as operational noise.

### 3.2. Change-Point Detection of Divergent Failure Modes

Running the change-point detection framework independently within each operational phase of the algal setup (System 1) identified fault times and degradation patterns across the reactors, successfully disaggregating physical structural failures from biological degradation, distinctions that aggregate mean-voltage plots obscure (Figure 2). During Phase 1 the algorithm flagged a single onset, in SMFC4 at *t* = 7.62 days, from which the reactor recovered. Its voltage fell by 8.0% to a minimum of 0.4991 V and returned to the level of the other bio-augmented reactors before the phase ended. No onset was detected in SMFC1, SMFC2, or SMFC3. During Phase 2, three of the four reactors registered onsets, at SMFC1 (*t* = 1.19 days), SMFC3 (*t* = 2.54 days), and SMFC4 (*t* = 4.33 days), with none in SMFC2. SMFC4 sustained by far the most severe collapse, a 64.3% voltage drop reaching a minimum of 0.1833 V, consistent with the physical separator damage. The non-algal control (SMFC3) showed an overall drop of 33.7% and the highest voltage fluctuation of the four reactors, reaching a peak rolling standard deviation of 0.1437 V at *t* = 7.9 days (a 7.1*×* increase over its baseline noise floor). The steep voltage collapse in SMFC4 reflects physical separator damage, while the highly fluctuating late-stage signal in SMFC3 indicates biological issues, such as biofilm detachment or mass-transfer limits at the anode. By isolating distinct failure modes, the change-point detection prevents mechanical faults from being misattributed to a lack of biocatalytic viability.

**Figure 2:**
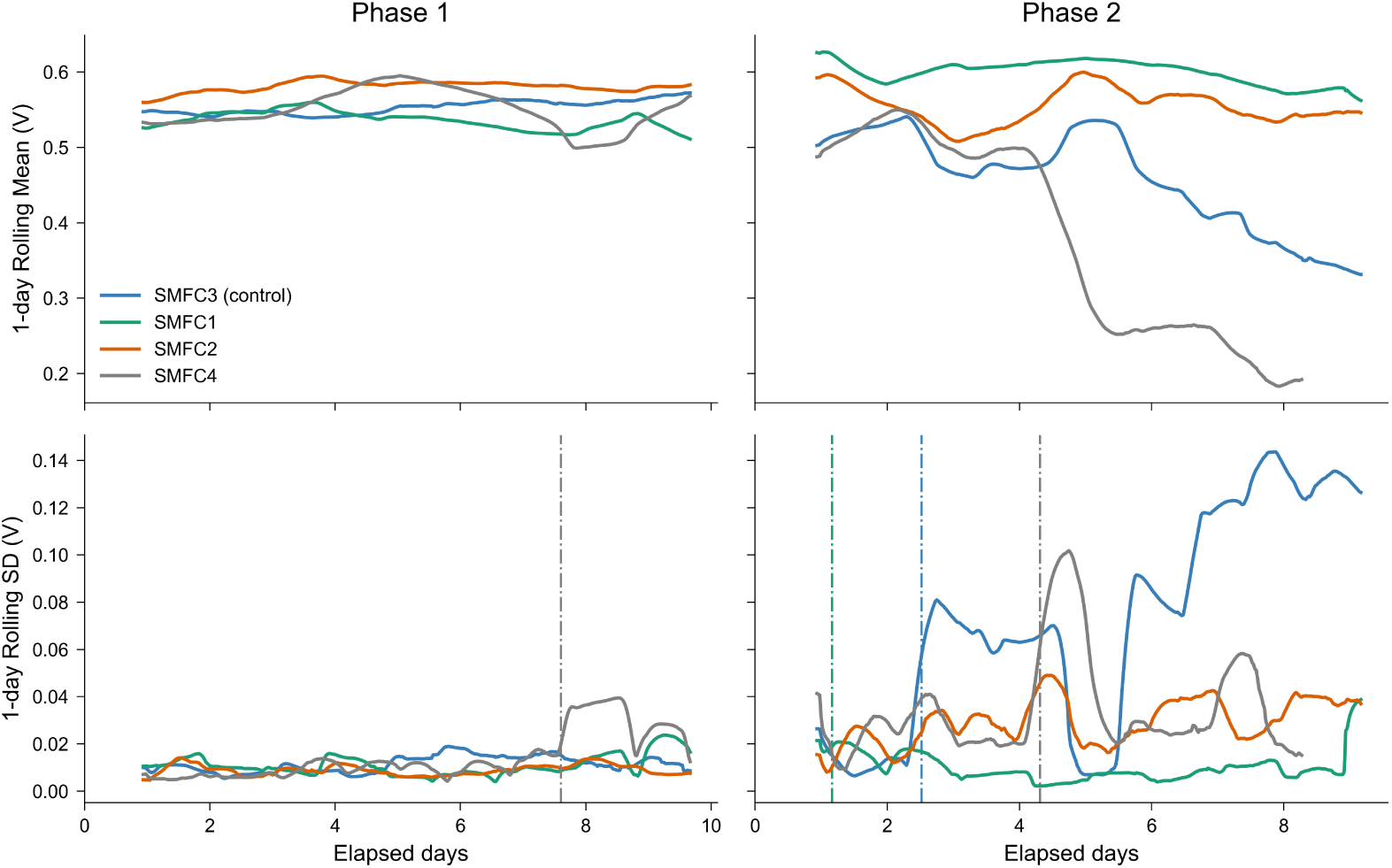
Change-point detection of reactor faults in System 1. One-day rolling mean voltage (top) and rolling standard deviation (bottom) for Phase 1 (left) and Phase 2 (right). Dashed vertical lines mark detected onsets.

To address cross-system generality, a prerequisite for scalable reactor health monitoring, the same framework was run separately on the secondary bacterial-consortia setup (System 2). The operation was performed without adjusting any algorithmic parameter other than the startup exclusion period, set to 6 days for this system against 1 day for System 1 (Figure 3). As with the first system, the framework successfully identified distinct fault patterns even though the setup had a different tailings substrate, sample rates, voltage ranges, and startup dynamics. In this system, the reactor T2 failed first (*t* = 6.27 days) with a minor voltage drop of 28.4%. T1 followed at *t* = 10.62 days with a substantial drop of 81.2%. Among the *P. alimentarius*-augmented reactors, C2 failed at *t* = 10.71 days (concurrent with T1) and C1 failed last at *t* = 12.81 days. Both exhibited the most severe voltage drops among all four reactors (C2: 108.8% and C1: 101.5%). C1 recorded the highest peak severity ratio (25.6*×* baseline) while C2 recorded the lowest (7.8*×*), reflecting its higher baseline noise floor (0.0169 V against 0.0090 V to 0.0099 V for the other three reactors). These drops included temporary dips below 0 V, which are characteristic of cell polarity reversal driven by anode bio-fouling or localized substrate exhaustion [30, 38]. Successfully deploying the framework, with only its startup exclusion adjusted, across two distinct systems confirms its reliability as an autonomous diagnostic tool for reactor performance. By detecting fault patterns consistent with both physical membrane damage and complex polarity reversals without manual tuning, the framework provides the necessary operational support for deploying bio-electrochemical reactors for continuous industrial wastewater management.

**Figure 3:**
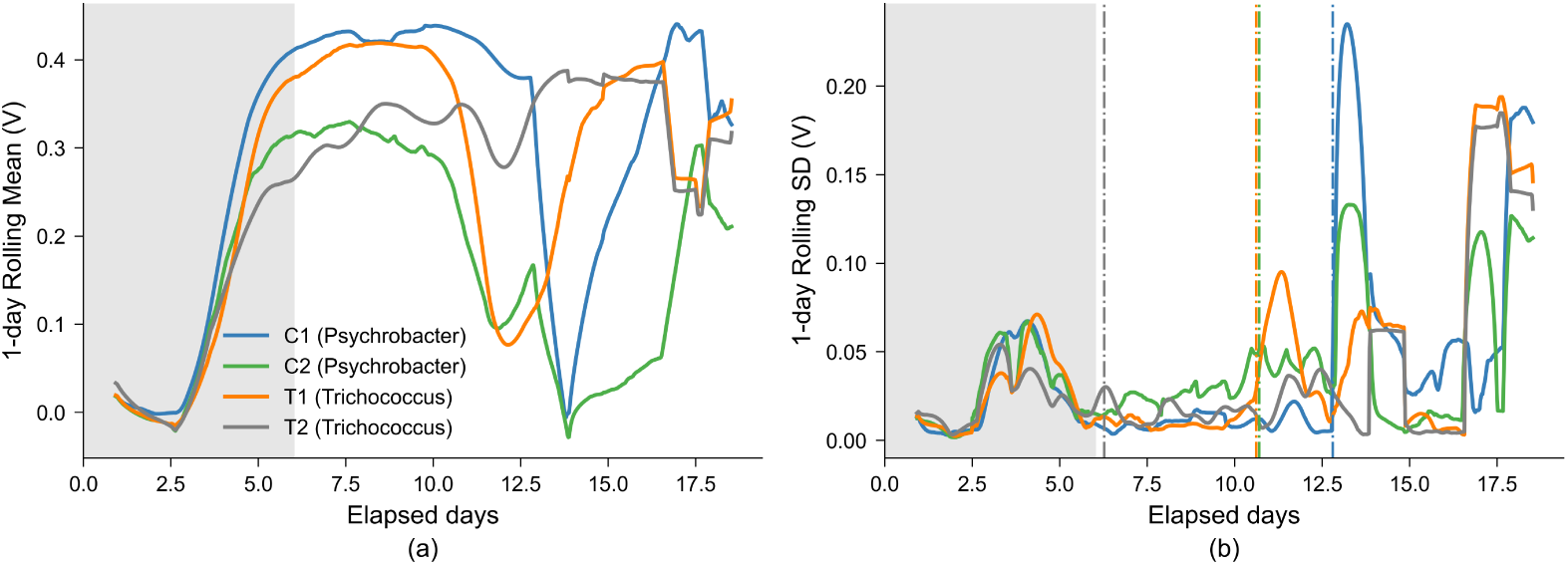
Change-point detection of reactor faults in System 2. Rolling mean voltage (a) and standard deviation (b) identify degradation onset (vertical lines). Shading marks 6-day startup exclusion. *Psychrobacter* reactors sustain deeper collapses and reverse polarity.

### 3.3. Diurnal Periodicity Explains Phase 2 Instability

We next evaluate the temporal patterns of the reactors in Phase 2. Figure 4 shows that the high variability in Phase 2 follows a daily cycle. Hour-of-day profiling reveals the non-algal control (SMFC3) oscillating with a peak-to-peak amplitude of 0.167 V, against 0.022 V to 0.057 V for the two algal reactors SMFC1 and SMFC2. Relative to day-to-day scatter, the control’s oscillation is 6.3 times its mean hourly standard error, compared with 5.1 for SMFC2 and 2.7 for SMFC1. For SMFC4, the apparent periodic structure is only 0.8 times its mean hourly standard error and cannot be distinguished from the collapse trend. The control reaches its minimum at 05:00 (*−*0.103 V relative to its phase mean) and its maximum at 18:00 (0.064 V), rising most steeply between 06:00 and 07:00 (0.084 V per hour). Lomb–Scargle analysis confirms the periodicity, showing that the dominant period is 24.3 h for the control (normalized power 0.380) and 24.2 h for SMFC2 (0.237), whereas SMFC1 shows no 24-hour component (dominant period 46.2 h) and SMFC4’s spectrum is dominated by its drop trend (56.3 h).

**Figure 4:**
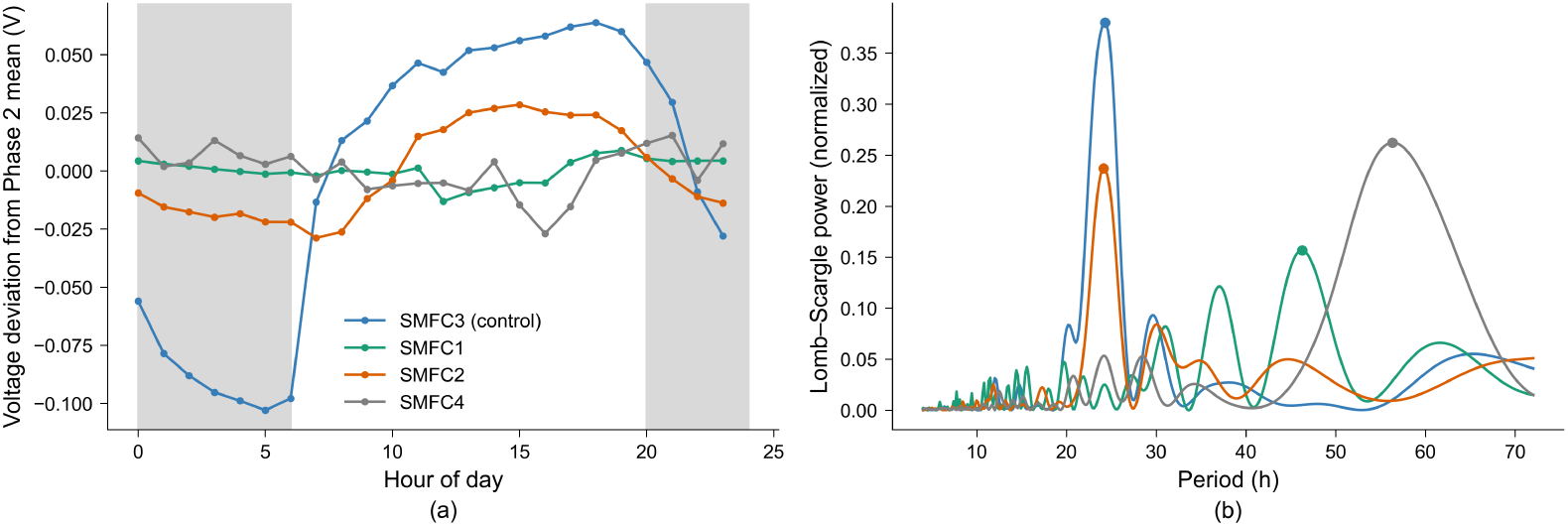
Phase 2 diurnal behaviour. (a) Hourly voltage deviations from phase means. (b) Lomb–Scargle periodograms identifying dominant periods. The control (SMFC3) shows the peak diurnal amplitude (0.167 V) and the strongest daily periodicity (24.3 h).

These transitions align with the chamber’s 06:00–20:00 photoperiod. Because all four reactors shared the same enclosure, temperature setpoint, illumination and physical positioning (Section 2.1), the differential response can be attributed to the reactors themselves rather than to their environment. These patterns indicate that in-situ photosynthetic oxygen evolution stabilizes the algal cathodes against diurnal cycling, whereas the passive-air control tracks these fluctuations directly. This provides a mechanistic account of the Phase 2 voltage and stability differences reported in Section 3.1. The results also demonstrate that periodicity analysis provides diagnostic information not captured by aggregate voltage statistics, as a reactor may maintain its mean operating voltage while developing a daily instability associated with reduced cathodic oxygenation.

### 3.4. Equivalent-Circuit Modeling of Interspecies Performance

Ohmic polarization modeling (Table 3, Figure 5) reveals that internal resistance (*R*_int_) in the reactors augmented with *T. patagoniensis* (T1, T2: 0.0325 kΩ *·* m^2^ to 0.0341 kΩ *·* m^2^) is approximately 40% lower than that of the reactors augmented with *P. alimentarius* (C1, C2: 0.0497 kΩ *·* m^2^ to 0.0601 kΩ *·* m^2^). The *T. patagoniensis* reactors also generated higher open-circuit voltages (OCV = 0.338 V to 0.386 V) than the *P. alimentarius* (OCV = 0.229 V to 0.258 V), consistent with reported differences in the electrogenic capacity of these taxa [27, 28]. The empirical maxima agree with Thévenin-predicted maximum power densities to within 30% of the observed value (Table 3), consistent with power performance being governed primarily by internal ohmic and thermodynamic polarization. By isolating these electrical parameters, the equivalent-circuit model successfully disaggregates the underlying bio-electrochemical kinetics from the background interference of the complex tailings.

**Figure 5:**
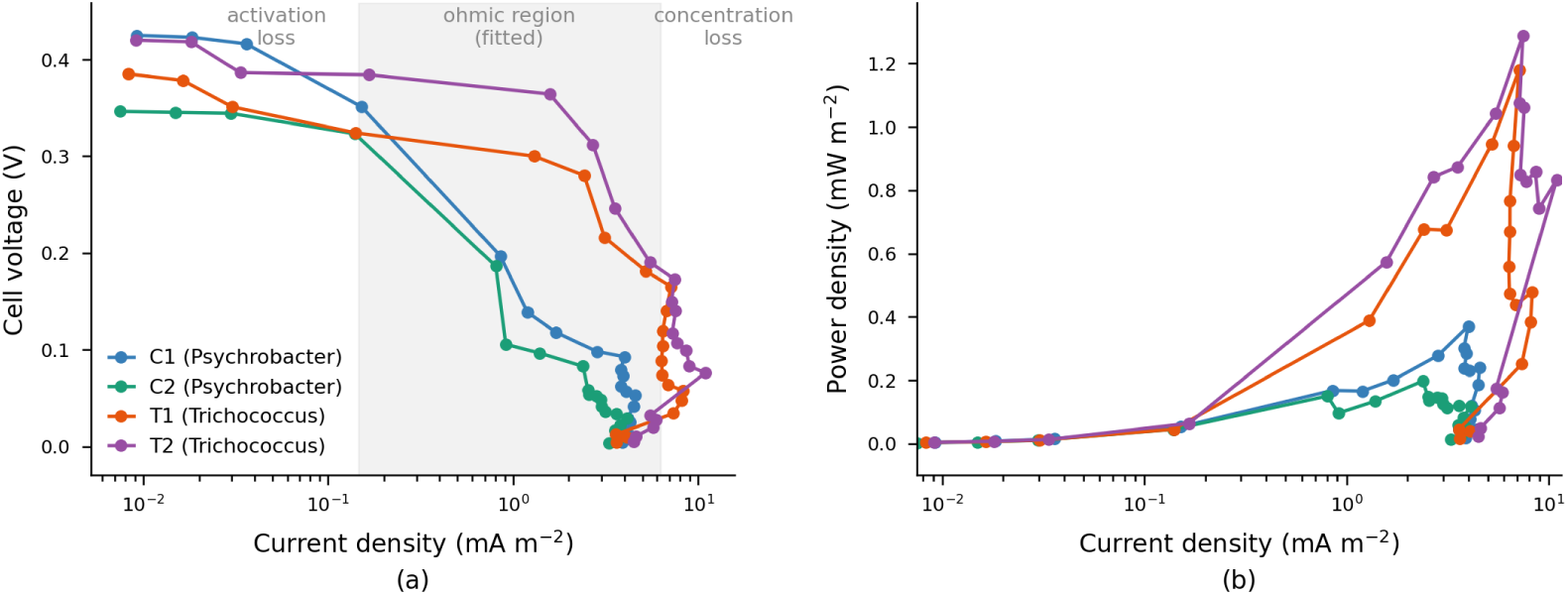
Interspecies polarization and power kinetics. Polarization (left) and power density curves (right) show roughly 4-fold higher peak power in *Trichococcus* (T1, T2) than *Psychrobacter* (C1, C2) due to higher open-circuit voltage and lower internal resistance.

**Table 3:** Equivalent-circuit fit parameters (ohmic region, *R*_ext_ = 300–100,000 Ω).

| Cell | OCV (V) | $R_{\text{int}}$ ($\text{k}\Omega\cdot\text{m}^2$) | $R^2$ | $P_{\text{max}}^{\text{obs}}$<br>($\text{mW}/\text{m}^2$) | $P_{\text{max}}^{\text{pred}}$<br>($\text{mW}/\text{m}^2$) |
| --- | --- | --- | --- | --- | --- |
| C1 (Psy.) | 0.258 | 0.0497 | 0.776 | 0.371 | 0.335 |
| C2 (Psy.) | 0.229 | 0.0601 | 0.739 | 0.197 | 0.217 |
| T1 (Tri.) | 0.338 | 0.0341 | 0.879 | 1.180 | 0.837 |
| T2 (Tri.) | 0.386 | 0.0325 | 0.948 | 1.289 | 1.145 |

Comparing internal resistance calculated from the voltage-current slope against estimates from the maximum power point (MPP) (Eq. 7, Table 4) highlights species-specific differences in transfer kinetics. For *T. patagoniensis* reactors, *R*_int_ estimates show strong agreement between the slope and MPP methods (T1: 0.0341 kΩ *·* m^2^ vs. 0.0308 kΩ *·* m^2^, *−*9.7%; T2: 0.0325 kΩ *·* m^2^ vs. 0.0331 kΩ *·* m^2^, +1.9%). Conversely, the estimates for the *P. alimentarius* reactors diverge (C1: +67%; C2: +84%). This discrepancy is linked to the operational voltage ratio (*V_P_*_max_ /OCV). The *T. patagoniensis* reactors operate close to the theoretical matched-load optimum (0.41–0.43 cf. ideal 0.50), maintaining ohmic polarization as current increases. In contrast, the *P. alimentarius* reactors operate far below this target (0.22–0.24), which causes non-ohmic polarization (resulting from activation overpotentials and mass-transport limitations near the peak power) to inflate the single-point MPP resistance estimates. This reconciliation remains a first-order proxy. Decoupling ohmic, charge-transfer, and mass-transport contributions requires electrochemical impedance spectroscopy. Leveraging this divergence between the resistance slope and MPP estimates provides a simple, non-invasive diagnostic check for identifying non-ohmic bottlenecks causing voltage loss. By helping to identify the kinetic limitations affecting reactor performance, the electrochemical modeling directly complements the statistical and computational analyses, completing the integrated diagnostic framework for continuous reactor monitoring.

**Table 4:** Reconciliation of slope-method vs. MPP-method internal resistance.

| Cell | $R_{\text{int}}^{\text{slope}}$<br>($\text{k}\Omega\cdot\text{m}^2$) | $R_{\text{int}}^{\text{MPP}}$<br>($\text{k}\Omega\cdot\text{m}^2$) | $V_{P_{\text{max}}}/\text{OCV}$ | Diff. (%) |
| --- | --- | --- | --- | --- |
| C1 (Psy.) | 0.0497 | 0.0831 | 0.218 | +67.2 |
| C2 (Psy.) | 0.0601 | 0.1106 | 0.239 | +84.1 |
| T1 (Tri.) | 0.0341 | 0.0308 | 0.429 | −9.7 |
| T2 (Tri.) | 0.0325 | 0.0331 | 0.411 | +1.9 |

### 3.5. Spectroscopic Characterization of Anode Surface Transformations

XRD profiles revealed clear mineral changes at the anode surface after treatment compared to the raw mine tailings (Figure 6). When normalized to the quartz (101) baseline, both treatments (SMFC-T and SMFC-C) exhibited strongly attenuated feldspar (2*θ* = 27.81*^∘^*) and calcite (2*θ* = 29.49*^∘^*) reflections. The relative intensity of feldspar decreased from 0.097 in the raw tailings to 0.064–0.066 in the treated anodes, while calcite intensity decreased from 0.122 to 0.078–0.082. These reductions are consistent with successful biological remediation of the tailings, further suggesting the effectiveness of the reactor designs. In parallel, the microbial breakdown of feldspar and calcite restructures the solid-liquid interface of the reactors, which relates to the internal-resistance differences indicated by the equivalent-circuit model. By physically confirming the dissolution of the mineral content, the XRD results suggest that the electrical modeling can accurately track dynamic in-situ remediation progress without requiring continuous physical sampling.

**Figure 6:**
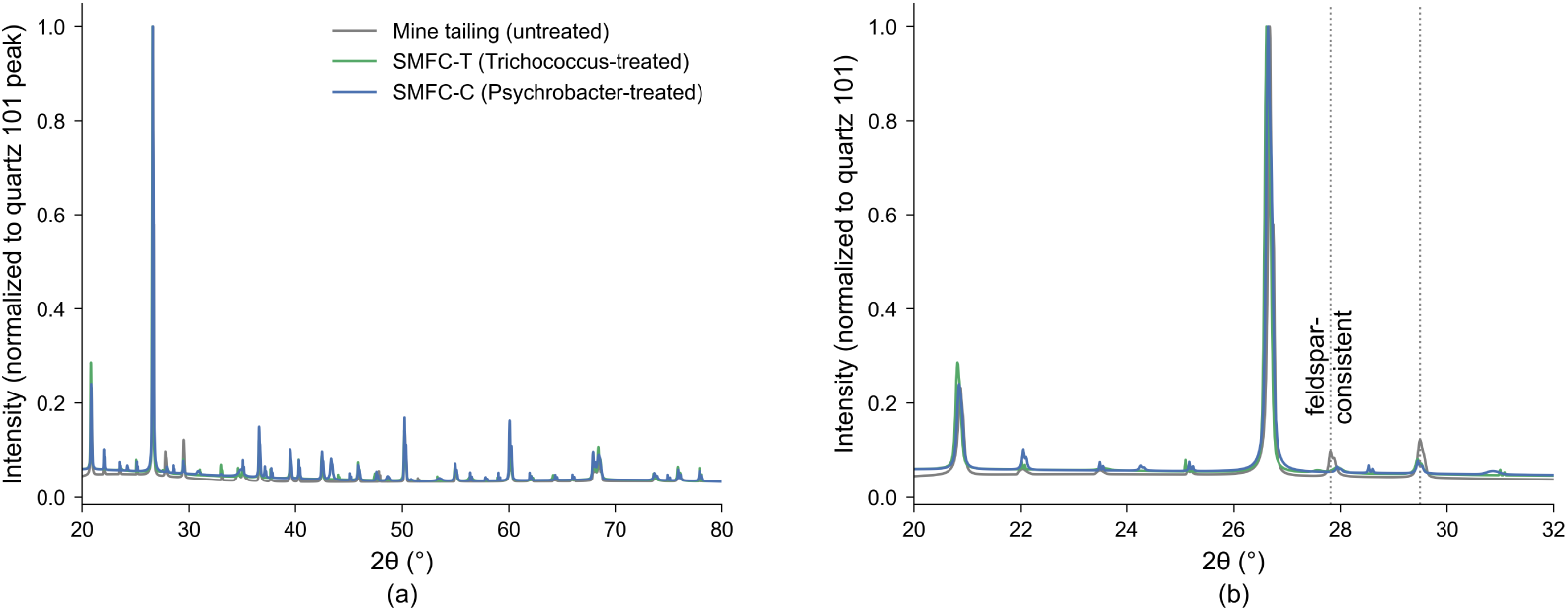
Tailings XRD mineralogy. (a) Quartz (101)-normalized patterns (20–32*^∘^* inset) (b) Zoom: Mineral phases beyond quartz. Feldspar and calcite attenuate relative to baseline, confirming biogenic dissolution.

FTIR spectroscopy indicated the development of a biogenic surface layer across all treated bio-anodes (Figure 7). Comparing the spectra revealed a stronger absorption band at *∼*1006 cm*^−^*^1^, which is characteristic of both polysaccharide C – O – C stretching in extracellular polymeric substances (EPS) and silica and carbonate materials. Treated anodes exhibited deeper peak absorption (16.1–17.5% below the baseline versus 12.9% in untreated tailings) and larger overall band areas (8.03–8.94 versus 6.17), as shown in Figure 7b. SEM observation of the bacterial biofilm morphology was consistent with these findings, indicating an EPS-rich biofilm on the electrode surface (Figure 7c). Because this biological layer simultaneously provides binding sites to immobilize metals and acts as a barrier that restricts mass transfer, its physical presence is consistent with the non-ohmic activation losses and diverging internal resistance estimates identified by the electrical modeling (Section 3.4) and suggests that the proposed integrated framework can help diagnose complex biological limitations in field-deployed reactors.

**Figure 7:**
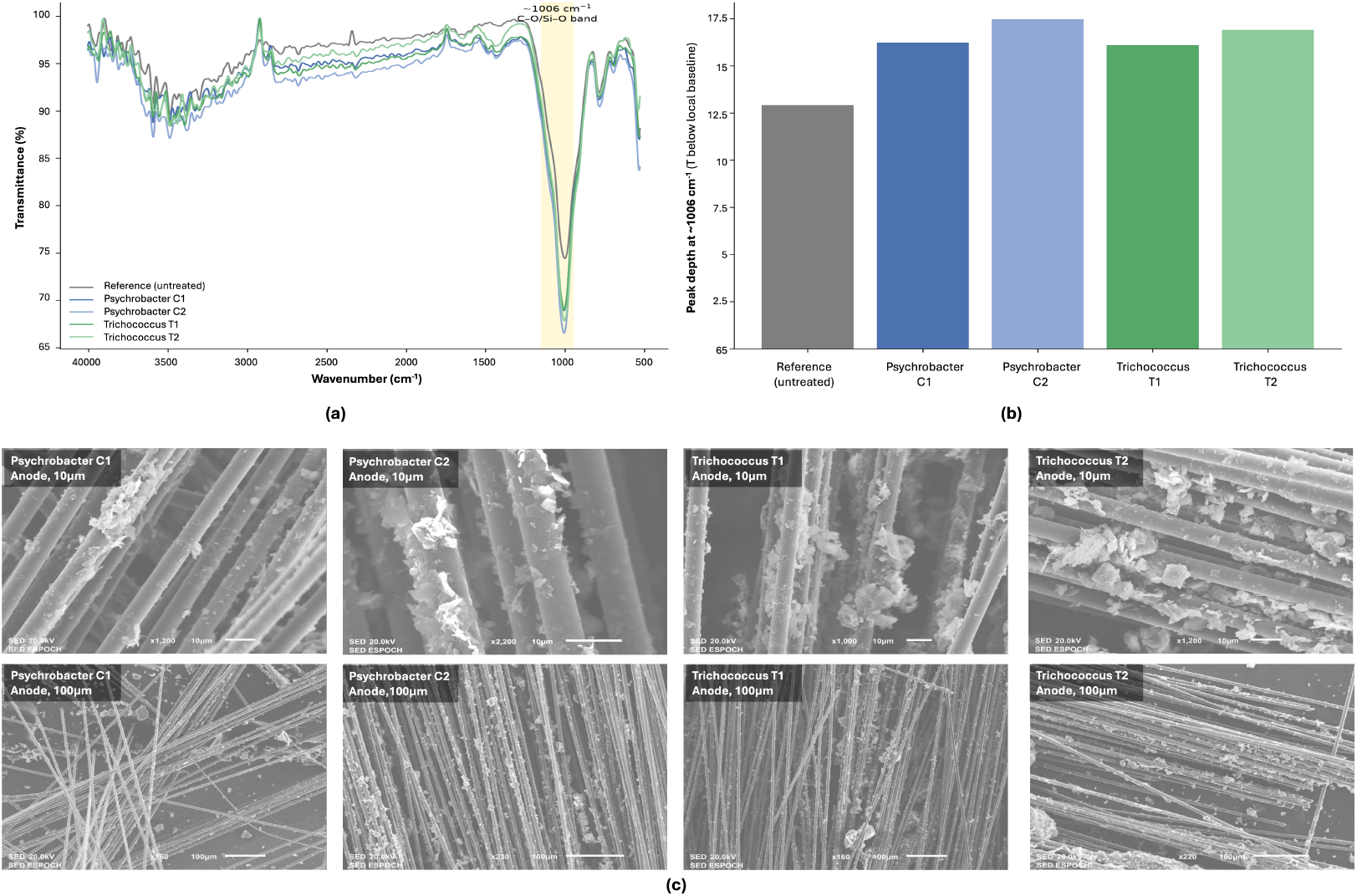
(a) FTIR spectra and *∼*1006 cm*^−^*^1^ band depth. (b) C-O/Si-O band strength. (c) Bacteria biofilm morphology by SEM. Increased absorption in treated samples vs. baseline confirms surface functionalization.

### 3.6. Systemic Bio-Electrochemical Drivers of Heavy Metal Remediation

Metal removal was quantified within each operational phase (Section 2.1). AAS analysis showed high depletion of copper and mercury in all reactors. The highest Phase 2 removals, Hg 93.5% and Cu 95.3%, were recorded in the non-algal control (SMFC3), but coincided with a severe operational trade-off characterized by an abrupt drop in voltage (Figure 8). In Phase 1 the ordering was reversed, with the algal reactors removing more Hg and Cu than the control. In the *M. inermum*-augmented units, mean removal was maintained from Phase 1 to Phase 2 for Cu (69.8% to 69.7%) and increased for Hg (81.4% to 90.5%). The average voltage in the algal units remained stable (0.56 V to 0.58 V), so remediation capacity was retained without the voltage loss recorded in the control.

**Figure 8:**
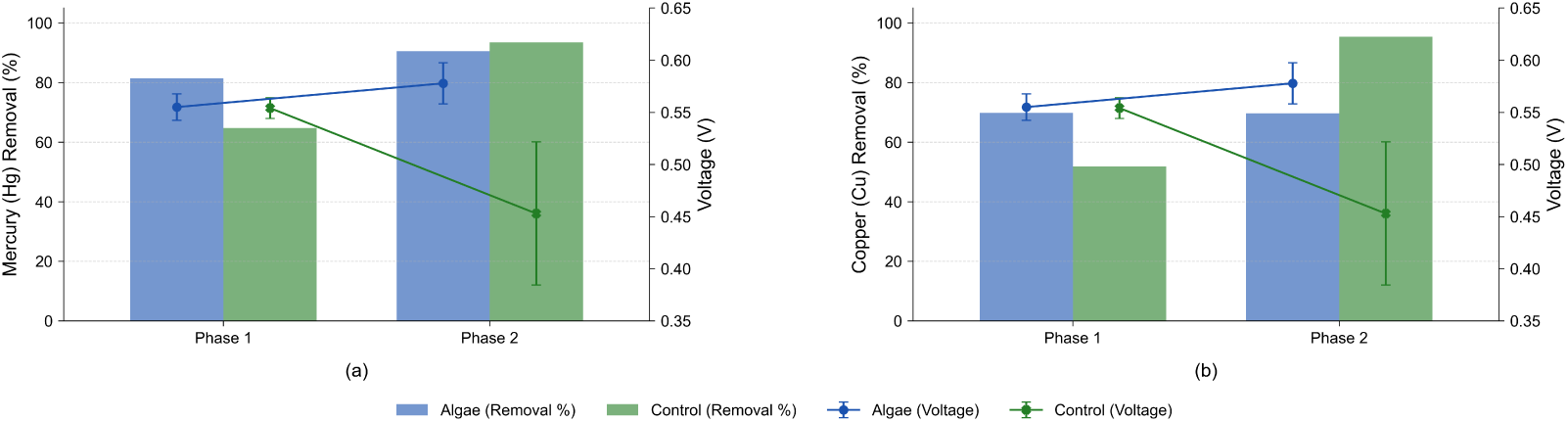
Trade-off between voltage and remediation: (a) Mercury, (b) Copper.

In terms of absolute concentrations, treatment reduced Hg from an influent of 0.042 mg L*^−^*^1^ to 0.0027–0.0148 mg L*^−^*^1^, and Cu from 2.79 mg L*^−^*^1^ to 0.130–1.837 mg L*^−^*^1^. Residual Pb was below the detection limit of the method in all reactors (Section 2.2). Benchmarked against the Ecuadorian discharge limits for freshwater bodies (Hg 0.001, Cu 1.0 mg L*^−^*^1^), Cu residuals complied in six of eight reactor–phase combinations, whereas Hg exceeded its limit in all eight. Clearing that threshold from the measured influent would require 97.6% Hg removal, above the 93.5% maximum observed here. Because regulatory thresholds for Hg are typically one to two orders of magnitude lower than those for Cu, Hg is the limiting contaminant in this matrix. While single-stage treatment in this configuration reduces metal loading, it falls short of discharge compliance for Hg.

The results demonstrate that substantial metal immobilization occurs independently of algal uptake and is instead associated with the shared bio-electrochemical environment. The EPS-rich surface layer indicated by FTIR and SEM provides binding sites for divalent cations (Cu^2+^, Hg^2+^), and micro-environmental gradients within the biofilm may drive precipitation of these species as hydroxide and carbonate phases. Because no external circuit was closed, these processes operate without faradaic electron transfer, which is consistent with removal being independent of the biocatalyst. These pathways are inferred: in the absence of a sterile abiotic control, passive adsorption onto the carbon electrodes, the separator and the mineral matrix cannot be distinguished from biologically mediated sorption and precipitation. Effluent pH fell from an influent value of 8.3 to 7.51–8.21 across all reactors and both phases (mean 7.82), so the reactors remained in the circumneutral to mildly alkaline range in which Cu^2+^ hydrolyses and is retained in the solid phase. This decrease in pH indicates a proton source exceeding the carbonate buffer, consistent with the partial calcite attenuation observed by XRD (Section 3.5). Because primary metal removal occurs independently of the algal population, biocatalyst selection can prioritize electrical stability without compromising remediation: an operator may adopt *M. inermum* to buffer against diurnal voltage loss while retaining the baseline metal recovery of the system. By mitigating the remediation–voltage trade-off, the algae-integrated system demonstrates a sustained capacity for dual-function operation, and the results further suggest that electrical fault detection can serve as an early-warning mechanism, isolating biological degradation prior to material loss in remediation capacity. The data nonetheless show that heavy-metal attenuation occurred across all reactors irrespective of algal augmentation.

### 3.7. Comparative Performance and Literature Benchmarking

Benchmarking reactor performance against existing algae- and consortium-based bio-electrochemical systems places the unbuffered, single-chamber mine-tailings architecture in context (Table 5). The peak power densities achieved by the bacterial-consortia system (System 2, 0.197 mW m*^−^*^2^ to 1.289 mW m*^−^*^2^) are consistent with values reported for systems employing mining waste as substrate (0.4 mW m*^−^*^2^ to 2.4 mW m*^−^*^2^) [47], and lower than those obtained on complex or synthetic wastewater (34 mW m*^−^*^2^ to 59 mW m*^−^*^2^) [24]. This gap is consistent with the electrical cost of the substrate. The fitted area-specific internal resistance is high (0.0325 kΩ·m^2^ to 0.0601 kΩ·m^2^) and maximum power occurs at external loads of 1 *−* 1.5 kΩ, reflecting the low conductivity and high overpotentials of raw, unbuffered mineral tailings. The *T. patagoniensis* cells recorded the highest values (1.180 mW m*^−^*^2^ to 1.289 mW m*^−^*^2^), and peak power varied 6.5-fold across the four cells..

**Table 5:** Comparison of this study’s mine-tailings MFC system with published bioelectrochemical / metal-removal MFC studies.

| Study | Organism | Substrate | Max. power (mW/m<sup>2</sup>) | Metal removal (%) | Key feature |
| --- | --- | --- | --- | --- | --- |
| [48] | Microalgae (cathodic) | Synthetic/lab medium | 0.128 mW total (not area-norm.) | not reported | Closed-cycle biomass + electricity co-generation |
| [22] | Microalgae-assisted, modified anode | Horse manure wastewater | 59 (best anode) | not reported | MWCNT/HNO<sub>3</sub> anode mod. raised power ~2.1–2.45× |
| [23] | *H. lacustris* (photosynthetic) | Synthetic catholyte | 34 (algae), 8 (control) | not reported | Algae-vs.-control design, analogous intent to this study’s System 1 |
| [21] | Various marine microalgae | Various | 345 (PMFC), 179 (MFC), 59 (BPV) mean | not reported | Cross-study means; power highly configuration-dependent |
| [12] | Mixed consortia | Synthetic/industrial | 400–6600 mW/m<sup>2</sup> | Cu: 99.9, Cr: 100, Cd: 90, Hg: 99.5, Ag/Au: 99.9 | Engineered two-chamber designs; near-quantitative removal |
| [49] | Mixed anaerobic/aerobic | Domestic/industrial | 45–5100 | not primary focus (COD 80–95) | Power strongly substrate- and configuration-dependent |
| **Ours** | *M. inermum*, *P. alimentarius* & *T. patagoniensis* | Gold-mine tailings | 0.197–1.289 (System 2), OCV up to 0.60 V (System 1) | Hg: 64.7–93.5, Cu: 34.2–95.3 | Paired reactor-day design separates conditional algae-specific voltage gain from large system-level metal removal |

Regarding heavy metal remediation, engineered two-chamber MFCs achieve near-quantitative recovery (> 99 %) [9], whereas the single-chamber algal design (System 1) achieves substantial attenuation directly within the untreated tailings matrix (Cu 34.2–95.3%, Hg 64.7–93.5%), with the non-algal control recording the highest Cu and Hg removals. The single-chamber configuration used substitutes a low-cost cellophane separator for a commercial ion-exchange membrane, and relies on system-level EPS bio-sorption and micro-environmental precipitation rather than membrane-mediated transport. Operationally, the electrical output of the consortium system and Phase 2 mercury attenuation of 86.5–93.1% in the algal system together establish a configuration that, paired with the proposed diagnostic framework, provides a baseline for automated health monitoring and process optimization in wastewater treatment.

### 3.8. Methodological scope and limitations

System 1 employed a single reactor per condition and one non-algal control. The paired reactor-day design addresses temporal pseudoreplication but does not substitute for biological replication. With only one control, between-reactor variance cannot be estimated independently of treatment. The Phase 1 spread among nominally identical algal reactors (*−*3.72% to +4.75% relative to the control) indicates that this variance is not negligible. The two phases were separated by physical reconstruction of all four cells, so the Phase 2 effect cannot be separated from the effect of the rebuild itself.

System 1 was operated under open-circuit conditions without an external load. The measured electrical quantity is cell potential rather than power, and metal removal cannot be attributed to current-driven reduction. No abiotic or sterile control was available in either system, so physicochemical sorption to the carbon electrode and mineral dissolution cannot be distinguished from biologically mediated uptake. The 0.15 V artefact-rejection threshold also excludes low readings from reactors approaching failure. Reported minimum voltages should be interpreted as upper bounds on the minimum voltage reached, while the corresponding magnitudes of voltage collapse represent lower bounds. XRD phase assignments were based on agreement with literature reference angles rather than full-pattern Rietveld refinement. Polarization-sweep data were available only for System 2, limiting the equivalent-circuit parameterization to that system. Removal efficiencies describe depletion of the dissolved phase, no solid-phase mass balance was performed.

## 4. Conclusions

This study establishes an integrated diagnostic and analytical framework for microbial fuel cells (MFCs) treating heavy-metal-rich gold mine tailings. Using a paired, per-day statistical design, the results show that while algal augmentation yields no detectable voltage advantage under stable conditions (Phase 1: +0.17%, *p* = 0.866), it drives a +27.54% enhancement (*p* = 0.0002) and stabilizes voltage under diurnal cycling. Heavy-metal remediation (up to 95.3%) is driven by shared system-level and physicochemical processes rather than algal-specific mechanisms. XRD and FTIR analyses indicate that the biological treatment attenuates key mineral phases and alters surface functional groups relative to the untreated tailings. An adaptive change-point algorithm successfully automated health monitoring across both independent test systems. It distinguished a recoverable voltage drop from a permanent physical collapse within the same reactor, while also flagging the non-algal control for severe late-stage instability. Equivalent-circuit modeling demonstrates that power density differences between microbial consortia result from combined shifts in internal resistance and open-circuit voltage. Together, these diagnostics provide a baseline for system monitoring, fault detection, and process optimization in bio-electrochemical tailings remediation and wastewater treatment. Future work should first establish whether metal removal is driven by physicochemical sorption or biologically mediated uptake through the inclusion of a sterile abiotic control and sequential extraction (BCR or Tessier) of the solid phases. Extending monitoring to continuous-flow operation, together with quantitative mineral tracking and time-resolved community sequencing, would establish whether exoelectrogens such as *T. patagoniensis* persist under sustained environmental stress. Techno-economic and life-cycle assessment is also needed to evaluate the viability of bio-augmented MFCs for industrial wastewater remediation.

## Supporting information

Algorithm

## CRediT Authorship Contribution Statement

**Joana Iza:** Conceptualization, Methodology, Formal analysis, Investigation, Writing (original draft). **Jennifer Cuadrado:** Conceptualization, Investigation, Validation. **Lourdes García-Rodríguez:** Supervision, Validation. **Celso Recalde:** Conceptualization, Project administration, Validation, Resources, Supervision. **Petteri Nurmi:** Methodology, Validation, Writing - review & editing. **Agustin Zuniga:** Conceptualization, Methodology, Software, Formal analysis, Validation, Writing - review & editing, Supervision.

## Data Availability

The 16S rRNA gene sequences of the bacterial isolates and the ITS region sequence of the microalga *M. inermum* will be deposited in NCBI GenBank upon acceptance. The change-point detection procedure is specified in full as pseudocode in Supplementary Information (Algorithm S1).

## Declarations

Ethics approval and consent to participate Ethical approval was not required for this study, as it did not involve human participants or live animals.

## Consent for publication

All authors have read and approved the final manuscript and consent to its publication.

## References

[1] L. Waaley, L. Mensah, S. Conrad, A. Gibrilla, S. Musah, T. H. Aiglsperger, G. K. Anornu, L. Alakangas, Environmental and health risk of artisanal and small-scale gold mining in Ghana: A review, Environmental Science and Pollution Research 33 (1) (2026) 1–16. 10.1007/s11356-025-37321-3.

[2] E. C. Nyanza, R. J. Mhana, M. Asori, D. S. Thomas, A. P. Kisoka, Effects of prenatal lead, mercury, cadmium, and arsenic exposure on children’s neurodevelopment in an artisanal small-scale gold mining area in northwestern tanzania using a multi-chemical exposure model, PLOS Global Public Health 5 (4) (2025) e0004577. 10.1371/journal.pgph.0004577.

[3] P. N. Obasi, B. B. Akudinobi, Potential health risk and levels of heavy metals in water resources of lead–zinc mining communities of abakaliki, southeast nigeria, Applied Water Science 10 (7) (2020) 184. 10.1007/s13201-020-01233-z.

[4] B. E. Logan, B. Hamelers, R. A. Rozendal, U. Schroder, J. Keller, S. Freguia, P. Aelterman, W. Verstraete, K. Rabaey, Microbial fuel cells: methodology and technology, Environmental Science and Technology 40 (17) (2006) 5181–5192. 10.1021/es0605016.

[5] K. Rabaey, W. Verstraete, Microbial fuel cells: novel biotechnology for energy generation, Trends in Biotechnology 23 (6) (2005) 291–298. 10.1016/j.tibtech.2005.04.008.

[6] B. E. Logan, Exoelectrogenic bacteria that power microbial fuel cells, Nature Reviews Microbiology 7 (5) (2009) 375–381. 10.1038/nrmicro2113.

[7] B. E. Logan, J. M. Regan, Electricity-producing bacterial communities in microbial fuel cells, Trends in Microbiology 14 (12) (2006) 512–518. 10.1016/j.tim.2006.10.003.

[8] L. S. Velez-Perez, J. Ramirez-Nava, G. Hern’andez-Flores, O. Talavera-Mendoza, C. Escamilla-Alvarado, H. M. Poggi-Varaldo, O. Solorza-Feria, J. A. Lopez-Diaz, Industrial acid mine drainage and municipal wastewater co-treatment by dual-chamber microbial fuel cells, International Journal of Hydrogen Energy 45 (25) (2020) 13720–13730. 10.1016/j.ijhydene.2019.12.037.

[9] A. T. Heijne, F. Liu, R. v. d. Weijden, J. Weijma, C. J. Buisman, H. V. Hamelers, Copper recovery combined with electricity production in a microbial fuel cell, Environmental Science and Technology 44 (11) (2010) 4376–4381. 10.1021/es100526g.

[10] H. Wang, X. Wang, Y. Zhang, D. Wang, X. Long, G. Chai, Z. Wang, H. Meng, C. Jiang, W. Dong, Y. Guo, J. Li, Z. Xu, Y. Lin, Sulfate-reducing bacteria-based bioelectrochemical system for heavy metal wastewater treatment: Mechanisms, operating factors, and future challenges, Journal of Electroanalytical Chemistry 952 (2023) 117967. 10.1016/j.jelechem.2023.117945.

[11] S. S. Lim, J.-M. Fontmorin, H. Pham, E. Milner, P. M. Abdul, K. Scott, I. Head, E. H. Yu, Zinc removal and recovery from industrial wastewater with a microbial fuel cell: Experimental investigation and theoretical prediction, Science of the Total Environment 776 (2021) 145979. 10.1016/j.scitotenv.2021.145934.

[12] S. Al-Asheh, M. Bagheri, A. Aidan, Removal of heavy metals from industrial wastewater using microbial fuel cell, Engineering in Life Sciences 22 (8) (2022) 535–549. 10.1002/elsc.202200009.

[13] J. V. Boas, V. B. Oliveira, M. Simões, A. M. Pinto, Review on microbial fuel cells applications, developments and costs, Journal of Environmental Management 307 (2022) 114525. 10.1016/j.jenvman.2022.114525.

[14] H. Bird, E. S. Heidrich, D. D. Leicester, P. Theodosiou, Pilot-scale microbial fuel cells (mfcs): a meta-analysis study to inform full-scale design principles for optimum wastewater treatment, Journal of Cleaner Production 346 (2022) 131227. 10.1016/j.jclepro.2022.131227.

[15] Y. Sun, H. Wang, X. Long, H. Xi, P. Biao, W. Yang, Advance in remediated of heavy metals by soil microbial fuel cells: Mechanism and application, Frontiers in Microbiology 13 (2022) 971589. 10.3389/fmicb.2022.997732.

[16] R. Rossi, B. E. Logan, Impact of reactor configuration on pilot-scale microbial fuel cell performance, Water Research 215 (2022) 118250. 10.1016/j.watres.2022.119179.

[17] Y. Dong, Z. Gao, J. Di, D. Wang, Z. Yang, X. Guo, X. Zhu, Study on the effectiveness of sulfate-reducing bacteria to remove Pb(II) and Zn(II) in tailings and acid mine drainage, Frontiers in Microbiology 15 (2024) 1369405. 10.3389/fmicb.2024.1352430.

[18] H. Wang, Y. Li, Y. Mi, D. Wang, Z. Wang, H. Meng, C. Jiang, W.-D. Dong, J. Li, H. Li, Cu(II) and Cr(VI) Removal in Tandem with Electricity Generation via Dual-Chamber Microbial Fuel Cells, Sustainability 15 (3) (2023) 2388. 10.3390/su15032388.

[19] E. E. Ziganshina, A. M. Ziganshin, Growth and productivity of micractinium inermum with increased inorganic carbon delivery under ammonium nutrition conditions, Phycology 6 (1) (2026) 26. 10.3390/phycology6010026.

[20] R. T. Smith, K. Bangert, S. J. Wilkinson, D. J. Gilmour, Synergistic carbon metabolism in a fast growing mixotrophic freshwater microalgal species micractinium inermum, Biomass and Bioenergy 82 (2015) 73–86. 10.1016/j.biombioe.2015.04.023.

[21] Z. H.-Y. Tay, F.-L. Ng, T.-C. Ling, M. Iwamoto, S.-M. Phang, The use of marine microalgae in microbial fuel cells, photosynthetic microbial fuel cells and biophotovoltaic platforms for bioelectricity generation, 3 Biotech 12 (7) (2022) 149. 10.1007/s13205-022-03214-2.

[22] N. Altın, B. Uyar, Increasing power generation and energy efficiency with modified anodes in algae-supported microbial fuel cells, Biomass Conversion and Biorefinery (2025). 10.1007/s13399-025-06536-2.

[23] A. Ahirwar, M. J. Khan, P. Khandelwal, G. Singh, Harish, V. Vinayak, M. M. Ghangrekar, Bioelectromics of a photosynthetic microalgae assisted microbial fuel cell for wastewater treatment and value added production, Scientific Reports (2025). 10.1038/s41598-025-13271-1.

[24] K. Obileke, H. Onyeaka, E. L. Meyer, N. Nwokolo, Microbial fuel cells, a renewable energy technology for bio-electricity generation: A mini-review, Electrochemistry Communications 125 (2021) 107003. 10.1016/j.elecom.2021.107003.

[25] D. D. Leicester, S. Settle, C. McCann, E. Heidrich, Investigating variability in microbial fuel cells, Applied and Environmental Microbiology 89 (3) (2023) e01662–22. 10.1128/aem.02181-22.

[26] L. Muhammad, Guidelines for repeated measures statistical analysis approaches with basic science research considerations, The Journal of Clinical Investigation 133 (11) (2023). 10.1172/JCI171058.

[27] K. Dai, Y. Yan, Q.-T. Wang, S.-J. Zheng, Z.-Q. Huang, T. Sun, R. J. Zeng, F. Zhang, Electricity production and key exoelectrogens in a mixed-culture psychrophilic microbial fuel cell at 4° c, Applied microbiology and biotechnology 106 (12) (2022) 4801–4811. 10.1007/s00253-022-12042-6.

[28] C. Armato, D. Ahmed, V. Agostino, D. Traversi, R. Degan, T. Tommasi, V. Margaria, A. Sacco, G. Gilli, M. Quaglio, et al., Anodic microbial community analysis of microbial fuel cells based on enriched inoculum from freshwater sediment, Bioprocess and biosystems engineering 42 (5) (2019) 697–709. 10.1007/s00449-019-02074-0.

[29] H. Wang, X. Long, Y. Sun, D. Wang, Z. Wang, H. Meng, C. Jiang, W. Dong, N. Lu, Electrochemical impedance spectroscopy applied to microbial fuel cells: A review, Frontiers in Microbiology 13 (2022) 973501. 10.3389/fmicb.2022.973501.

[30] B. Kim, I. S. Chang, R. Dinsdale, A. Guwy, Accurate measurement of internal resistance in microbial fuel cells by improved scanning electrochemical impedance spectroscopy, Electrochimica Acta 366 (2021) 137388. 10.1016/j.electacta.2020.137388.

[31] S. P. Jung, S. Son, B. Koo, Reproducible polarization test methods and fair evaluation of polarization data by using interconversion factors in a single chamber cubic microbial fuel cell with a brush anode, Journal of Cleaner Production 390 (2023) 136157. 10.1016/j.jclepro.2023.136157.

[32] W. Taufemback, D. Hotza, D. Recouvreux, P. C. Calegari, T. Pineda-Vásquez, R. Antônio, E. Watzko, Techniques for obtaining and mathematical modeling of polarization curves in microbial fuel cells, Materials Chemistry and Physics (2024). 10.1016/j.matchemphys.2024.128998.

[33] J. Greenman, I. Gajda, J. You, A. Mendis, O. Obata, G. Pasternak, I. Ieropoulos, Microbial fuel cells and their electrified biofilms, Biofilm 3 (2021). 10.1016/j.bioflm.2021.100057.

[34] I. M. Simeon, A. Weig, R. Freitag, Optimization of soil microbial fuel cell for sustainable bio-electricity production: combined effects of electrode material, electrode spacing, and substrate feeding frequency on power generation and microbial community diversity, Biotechnology for Biofuels and Bioproducts 15 (2022). 10.1186/s13068-022-02224-9.

[35] T. Kamperidis, P. Pandis, C. Argirusis, G. Lyberatos, A. Tremouli, Effect of Food Waste Condensate Concentration on the Performance of Microbial Fuel Cells with Different Cathode Assemblies, Sustainability (2022). 10.3390/su14052625.

[36] M. Blatter, L. Delabays, C. Furrer, G. Huguenin, C. Cachelin, F. Fischer, Stretched 1000-l microbial fuel cell, Journal of Power Sources 483 (2021) 229148. 10.1016/j.jpowsour.2020.229130.

[37] P. Jalili, A. Ala, P. Nazari, B. Jalili, D. D. Ganji, A comprehensive review of microbial fuel cells considering materials, methods, structures, and microorganisms, Heliyon 10 (3) (2024) e24804. 10.1016/j.heliyon.2024.e25439.

[38] F. Fischer, N. Merino, M. Sugnaux, G. Huguenin, K. Nealson, Microbial community diversity changes during voltage reversal repair in a 12-unit microbial fuel cell, Chemical Engineering Journal (2022). 10.1016/j.cej.2022.137334.

[39] S. Schmidl, P. Wenig, T. Papenbrock, Anomaly detection in time series: A comprehensive evaluation, Proceedings of the VLDB Endowment 15 (9) (2022) 1779–1797. 10.14778/3538598.3538602.

[40] P. Du, N. M. Abdel-Jabbar, B. A. Wilhite, C. Kravaris, Fault Diagnosis in Chemical Reactors with Data-Driven Methods, Industrial and Engineering Chemistry Research 64 (11) (2025) 6060–6076. 10.1021/acs.iecr.4c04042.

[41] D. Fernández-Verdejo, P. Cortés, A. Guisasola, P. Blánquez, E. Marco-Urrea, Bioelectrochemically-assisted degradation of chloroform by a co-culture of *Dehalobacter* and *Dehalobacterium*, Environmental Science and Ecotechnology 12 (2022). 10.1016/j.ese.2022.100199.

[42] W. Kong, Y. Li, Y. Zhang, H. Liu, Enhanced degradation of refractory organics by bioelectrochemical systems: A review, Journal of Cleaner Production (2023). 10.1016/j.jclepro.2023.138675.

[43] Ministerio del Ambiente del Ecuador, Acuerdo Ministerial 097-A: Anexo 1 del Libro VI del Texto Unificado de Legislación Secundaria del Ministerio del Ambiente, Norma de Calidad Ambiental y de Descarga de Efluentes al Recurso Agua, Registro Oficial, Edición Especial No. 387, published 4 November 2015 (Nov. 2015). URL https://www.gob.ec/sites/default/files/regulations/2018-09/Documento_Acuerdo-Ministerial-097-A-ANEXOS.pdf

[44] A. Erensoy, N. Çek, Investigation of polymer biofilm formation on titanium-based anode surface in microbial fuel cells with poplar substrate, Polymers 13 (2021). 10.3390/polym13111833.

[45] T. Eyoel, Y. Shuka, S. Tadesse, T. Tesfaye, M. Mengesha, Green Energy: Power Generation Improvement in Microbial Fuel Cells Using Bio-Synthesized Polyaniline-Coated Co_3_O_4_ Nanocomposite, International Journal of Energy Research 2025 (2025). 10.1155/er/2936572.

[46] A. Adekunle, C. Rickwood, B. Tartakovsky, Online monitoring of heavy metal–related toxicity using flow-through and floating microbial fuel cell biosensors, Environmental Monitoring and Assessment 192 (1) (2020) 52. 10.1007/s10661-019-7850-0.

[47] E. Leiva-Aravena, E. Leiva, V. Zamorano, C. Rojas, J. M. Regan, I. T. Vargas, Organotrophic acid-tolerant microorganisms enriched from an acid mine drainage affected environment as inoculum for microbial fuel cells, Science of the Total Environment 678 (2019) 639–646. 10.1016/j.scitotenv.2019.05.003.

[48] I. Gajda, J. Greenman, I. A. Ieropoulos, Recent advancements in real-world microbial fuel cell applications, Current opinion in electrochemistry 11 (2018) 78–83. 10.1016/j.coelec.2018.09.006.

[49] H. Roy, T. U. Rahman, N. Tasnim, J. Arju, M. M. Rafid, M. R. Islam, M. N. Pervez, Y. Cai, V. Naddeo, M. S. Islam, Microbial Fuel Cell Construction Features and Application for Sustainable Wastewater Treatment, Membranes 13 (5) (2023) 490. 10.3390/membranes13050490.

