## Supplementary material for "Removal of Hg and Cu from raw gold-mine tailings in microalgae- and bacteria-augmented microbial fuel cells: Effects of bio-augmentation, diurnal cycling and operational faults": Algorithm

[a]*Escuela Superior Politecnica del Chimborazo, Avenida Pedro Vicente Maldonado s/n y Avenida 11 de Noviembre, Riobamba, 060155, Chimborazo, Ecuador*
[b]*Departamento de Ingeniería Energética, Escuela Técnica Superior de Ingeniería (ETSI), Universidad de Sevilla, ETSI, Camino de los Descubrimientos, s/n, Seville, 41092, Spain*
[c]*ENGREEN, Laboratorio de Ingeniería para la Sostenibilidad Energética y Medioambiental – Unidad de Excelencia de la Universidad de Sevilla, ETSI, Camino de los Descubrimientos, s/n, Seville, 41092, Spain*
[d]*University of Helsinki, P.O. Box 68 (Pietari Kalmin katu 5), Helsinki, FI-00014 University of Helsinki, Uusimaa, Finland*

### S1. Algorithm

Algorithm S1 gives the pseudocode for the adaptive percentile-floor change-point detector described in Section 2.3. Symbols follow the main-text definitions: $v(t)$ is the raw voltage series, $W$ the 1-day rolling window (48 intervals of 30 minutes), $\sigma(t)$ the rolling standard deviation, $F$ the per-reactor noise floor, and $t_{\text{exclude}}$ the startup exclusion time, set to 1 day for each phase of System 1 and 6 days for System 2.

*Corresponding author.

**Algorithm S1:** Percentile-floor change-point detection and degradation ranking algorithm.

**Input:** Voltage series $v(t)$, rolling window $W$, startup exclusion time $t_{\text{exclude}}$, phase boundaries

**Output:** Per phase: degradation onset time $t_0$, severity ratio $S$, relative voltage collapse $D$, reactor rank matrix $(t_0, \sigma_{\max}, S)$

**foreach** *operational phase* **do**
- Reference $t$ to the phase start;
- Resample $v(t)$ to 30-minute interval means and standard deviations;
- Exclude windows with incomplete coverage;
- Compute rolling standard deviation $\sigma(t)$ over moving window $W$;
- Compute baseline noise floor $F$ via Eq. (2);
- **foreach** *reactor* **do**
  - **for** $t \geq t_{exclude}$ *in chronological order* **do**
    - **if** $\sigma(t) \geq 3F$ *sustained for* $\geq 70\%$ *of the subsequent 6 intervals (Eq. 3)* **then**
      - $t_0 \leftarrow t$;
      - **break**;
    - **end**
  - **end**
  - $\sigma_{\max} \leftarrow \max_t \sigma(t)$;
  - $S \leftarrow \sigma_{\max}/F$;
  - Compute voltage collapse $D$ via Eq. (5);
- **end**
- Rank reactors sequentially by primary criterion $t_0$, secondary criterion $\sigma_{\max}$, and tertiary criterion $S$;

**end**
